# Delineating the effects of prenatal oxycodone exposure and melatonin treatment on placental and fetal outcomes in pregnant rats

**DOI:** 10.64898/2026.04.23.720463

**Authors:** IO Adediji, HM Kowash, P Nouri Mousa, CO Aloba, VL Schaal, JS Davis, ES Peeples, G Pendyala, LK Harris

## Abstract

**Background:** Prenatal oxycodone (oxy) exposure has been associated with adverse pregnancy and fetal developmental outcomes. In this study, we assessed whether chronic prenatal oxy exposure impairs placental and fetal growth in rats and if maternal melatonin supplementation would mitigate these effects.

**Methods:** Female Sprague-Dawley rats received either saline or oxy via oral gavage for 15 days before mating (10-15mg/kg dose escalation) and throughout pregnancy (15mg/kg). From gestational day (GD) 12.5, half of the dams received melatonin (10mg/kg). On GD19.5, maternal and fetal blood, and maternal, placental and fetal tissues were harvested. Placental histomorphometry was assessed and immunohistochemistry for pan-cytokeratin, PCNA, CD34, α-SMA, and TUNEL analysis were performed. Maternal and fetal plasma cytokines, angiogenic factors, and pregnancy hormones were measured by ELISA. Anthropometric data were analyzed using general linear mixed models and other outcomes were analyzed using univariate general linear models.

**Results:** Oxy induced fetal growth restriction as evidenced by reduced placental weight, fetal weight, fetal-to-placental weight ratio, crown-rump length, and fetal liver weight. Melatonin also independently reduced some parameters of fetal growth but when administered with oxy it partially improved fetal outcomes including the head-to-abdominal diameter ratio. Oxy exposure increased placental labyrinth zone area, the percentage of CD34-positive cells, and maternal plasma IL-1β and IL-10 concentrations and reduced the percentage of pan-cytokeratin positive cells, while both oxy and melatonin reduced maternal plasma chorionic gonadotropin levels.

**Conclusion:** Prenatal oxy exposure disrupts placental structure, labyrinth anatomy, and induces maternal systemic inflammation, associated with impaired fetal growth. The protective effects of melatonin are partial but indicate a potential brain sparing effect.

## INTRODUCTION

The widespread misuse of prescription opioids and the increased availability of illicit opioids has contributed to what is now widely recognized as an opioid epidemic^1^. Use of opioids such as morphine, methadone and oxycodone (oxy) during pregnancy has increased rapidly in recent years, which poses significant risks to the developing fetus, as many opioids cross the placental barrier^2–4^. Between 2008-2012, the Center for Disease Control and Prevention (CDC) reported that a third of reproductive aged women receiving Medicaid support, and over 25% of women using private healthcare insurance had at least one opioid prescription filled annually^5^. From 2010 to 2017, the estimated rate of maternal opioid-use disorder (MOUD) diagnoses increased from 3.5 per 1000 hospital deliveries, to 8.2 per 1000 hospital deliveries in the US, with significant variation between states. In parallel, the estimated rate of neonatal abstinence syndrome (NOWS) increased from 4.0 to 7.3 per 1000 hospital deliveries^6^. Maternal opioid use disorder and chronic opioid exposure during pregnancy are associated with a range of adverse maternal, placental and fetal outcomes, including alterations in placental structure and function^7^, and increased incidence of fetal growth restriction (FGR), preterm birth and stillbirth^8–10^ Furthermore, opioid exposure in utero is associated with detrimental neurodevelopmental outcomes later in life including impaired cognition, motor skills and sensory processing, and behavioral deficits^11–14^.

The placenta is a central regulator of fetal growth and development, acting via a diverse range of anatomical, physiological and endocrine mechanisms^15, 16^. In early pregnancy, remodeling of uterine spiral arteries by a subset of invasive extravillous trophoblast cells ensures delivery of sufficient blood to maintain adequate placental perfusion^17^. Rapid growth of the placental villous tissue provides a large surface area equipped with specialized transporters to facilitate nutrient and gas exchange^18^. Additionally, the placenta is a critical mediator of endocrine and immune function via the production and release of hormones, cytokines and chemokines^18^. Disruption of these tightly regulated processes can impair placental growth, development and uteroplacental perfusion^19^, leading to a host of cellular and molecular changes resulting in reduced nutrient transport capacity^18^, oxidative damage and trophoblast mitochondrial dysfunction^20^, inflammation and apoptosis^17^. Collectively, these insults can disrupt fetal organ development, impair fetal growth trajectory and lead to developmental programming of long-term disease in affected offspring^18, 21, 22^.

There have been limited studies to date on the effects of MOUD on human placental structure and function but delayed maturation of placental villi and placentomegaly have been reported^23, 24^. The κ- and μ-opioid receptors are expressed by human villous trophoblast lineages^7^ and opioids are known to cause vasoconstriction of placental vessels and release of human chorionic gonadotropin (hCG) and human placental lactogen (hPL)^7, 25–27^. Given that oxy has the highest rate of placental transfer compared to morphine, heroin, fentanyl and methadone^28^ and unlike these other opioids, no efflux transporters exist that can actively remove oxy from placental tissue^29, 30^ there is therefore increasing concern regarding the potential impact of oxy on placental function, fetal development and pregnancy outcomes.

We have previously established a rat model of prenatal oxy exposure which recapitulates the fetal growth restriction observed in human infants^31, 32^. This model also exhibits abnormalities in fetal and neonatal brain development, including increased concentrations of the neurotransmitter glutamate, and changes in the expression of genes related to inflammation, opioid signaling, pain perception, learning and memory^31–37^ Collectively, these findings demonstrate that oxy exposure disrupts both developmental and neurobiological trajectories, but the underlying mechanisms remain unclear and require further investigation. Additionally, appropriate interventions to mitigate maternal pregnancy complication and fetal brain injury during oxy exposure are lacking.

Melatonin is an endogenous indoleamine hormone produced primarily by the pineal gland but also by the placenta. It is typically known for its role in regulation of circadian rhythms^38, 39^, but it also has antioxidant, anti-apoptotic, anti-inflammatory and neuroprotective properties^40^. Compelling evidence from a number of pre-clinical pregnancy models indicates that melatonin can reduce placental oxidative stress and inflammation^40^ improve uteroplacental vascular flow and reduce placental ischemia^41^ promote fetal growth and overall survival^42^ and reduce inflammation and exert neuroprotective effects on the fetal brain^42, 43^ Given that melatonin is deemed safe for use in pregnancy and is currently being evaluated in clinical trials to prevent fetal brain injury in FGR-affected pregnancies (NCT01695070)^44, 45^ and mitigate brain impairment caused by premature delivery (NCT04235673)^46^, we sought to assess its ability to protect against the adverse effects of oxy exposure. Accordingly, we hypothesized that prolonged maternal oxy exposure in a rat model leads to alterations in maternal pregnancy physiology, placental development, and fetal outcomes and that the administration of melatonin could mitigate these opioid-induced changes.

## MATERIALS AND METHODS

### Animals and Experimental Design

All animal protocols were approved by the Institutional Animal Care and Use Committee of the University of Nebraska Medical Center (UNMC), and were conducted in accordance with the National Institutes of Health Guide for the Care and Use of Laboratory Animals^47^. Male and female Sprague Dawley rats were obtained from Charles River Laboratories Inc. (Wilmington, MA, USA), and group housed in a 12-h light–dark cycle and fed ad libitum. Oxycodone HCl (Sigma Aldrich, St. Louis, MO, U.S.A.) exposure was adapted from previously published studies^31, 34, 48^. Melatonin treatment was performed in accordance with previously published work^42, 43, 49^. Briefly, between 8 am and 9 am each morning, nulliparous female (64-70 days of age) Sprague-Dawley rats received either saline or oxy by oral gavage for 5 days, starting with a dose of 10 mg/kg/day, which was then increased by 0.5 mg/kg/day over the next 10 days to reach a final dose of 15 mg/kg/day. On day 15, the female rats were mated with proven male breeders. Pregnancy was confirmed by the presence of a vaginal plug the following morning, and the day of plugging was designated as gestational day (GD) 0.5. This treatment regimen continued throughout gestation. From GD12.5 to GD19.5, half of the dams in each group additionally received oral melatonin (Sigma Chemical Co., St. Louis, MO, USA) at a dose of 10 mg/kg, resulting in four experimental groups: saline (Sal), saline and melatonin (Mel), Oxy, and Oxy and melatonin (OxMe) (Supplementary Figure S1).

### Tissue Harvest

On GD19.5, dams were euthanized by isoflurane overdose (5%; Abbott, UK) one hour after the final treatment. Maternal blood was collected by cardiac puncture and centrifuged at 1000 x g at 4°C for 10 minutes; plasma was aliquoted into sterile tubes and snap frozen. The uterine horn was removed and placed on ice; fetuses and placentas were removed and weighed. Maternal and fetal organs were then dissected, weighed and either snap frozen, stored in RNA later or fixed for histological examination. Trunk blood was collected from each fetus and pooled within a litter to obtain sufficient plasma volume for cytokine quantification. Fetal tail tips were collected for sex determination, performed by PCR amplification of the male-specific Sry gene, as previously described^50, 51^.

### Placental histomorphometry

Following harvest, placental tissue was fixed in 10% (v/v) neutral buffered formalin for 24h, washed in PBS and paraffin-embedded. Sections (5 µm) were cut using a rotary microtome (Leica RM2125 RTS; Leica Biosystems, UK) and stained with hematoxylin (Stat Lab #95-1) and eosin (Stat Lab #98-1), as previously described^52^. Three sections per slide were imaged by brightfield microscopy using the Aperio GT Elite slide scanner. Placental junctional zone and labyrinth zones were manually demarcated using the using the freehand tool in Image J (NIH, USA) and the areas were quantified at 1X magnification using the software as previously described^53^.

### Immunohistochemistry

Paraffin-embedded placenta sections were dewaxed and rehydrated in descending grades of alcohol, followed by heat-induced antigen retrieval in 0.01M sodium citrate buffer (pH 6.0; Sigma-Aldrich) for 10 minutes. After cooling, endogenous peroxidase activity was quenched with 3% (w/v) hydrogen peroxide in water (Fisher #7722-84-1) for 10 minutes. Sections were washed three times in 1X TBS containing 1% (v/v) Tween-20 (1X TBST; 5 minutes per wash) and blocked with 10% (v/v) normal goat serum (Abcam #7481) in 1X TBST for 30 minutes at room temperature. Primary antibodies were diluted in 10% normal goat serum and incubation was carried out overnight at 4°C using the following dilutions: anti-pan cytokeratin (1:200; mouse mAb #sc-8018, Santa Cruz Biotechnology), anti-PCNA (1:100; rabbit mAb #5701035, Sigma-Aldrich), anti-αSMA (1:500; rabbit mAb #30894, Novus Biotechne), and anti-CD34 (1:500; Abcam, rabbit mAb #316277). Sections processed without primary antibody served as negative controls. Following overnight incubation, sections were washed in 1X TBST (0.6% Tween-20) and incubated for one hour with appropriate secondary antibodies (goat anti-mouse; Vector Lab #BA-9200 or goat anti-rabbit; Vector Lab #BA-1000). After three additional washes with 1X TBST (0.6% Tween-20), sections were incubated with avidin-peroxidase (1:200; Vector Laboratories #A-2004) for 30 minutes. Signal was developed using 3,3’-diaminobenzidine (Sigma-Aldrich #D4293) for 1 to 2 minutes, followed by rinsing in water, counterstaining with hematoxylin (Stat Lab #95-1), dehydration, and mounted using CoverSafe mounting medium (Stat Lab #MMC0226). Cell death was assessed in parallel using a TUNEL assay kit performed, per the manufacturer’s protocol (Abcam #ab206386). Stained slides were scanned by brightfield microscopy using the Aperio GT Elite slide scanner at the Tissue Science Facility, UNMC, Omaha. Ten fields per placental zone (labyrinth and junctional) were analyzed per placenta, and the number of positively stained cells were quantified using QuPath version 0.1.5 (University of Edinburgh; https://qupath.github.io).

### ELISA

Maternal and fetal plasma cytokine concentrations (IL-1β, IL-6, TNF-α, CXCL1, IL-2, IL-10, IL-4, and IL-5) were quantified in duplicates using the V-PLEX Proinflammatory Panel 1 Rat Kit (K15294G, Meso Scale Discovery, Rockville, MD), according to the manufacturer’s instructions. Briefly, plasma samples and calibrators were added to pre-coated multi-spot plates and incubated to allow analyte binding. Following washing, detection antibodies conjugated with electrochemiluminescent labels were applied. After a final wash, read buffer was added, and plates were analyzed using a MESO QuickPlex SQ 120 instrument. Signals were quantified via electrochemiluminescence, and cytokine concentrations were calculated from standard curves generated for each analyte. The concentrations of placenta growth factor (PlGF; #MBS026910), vascular endothelial growth factor (VEGF; #MBS161464), soluble fms-like tyrosine kinase 1 (sFlt-1; #MBS2602003), chorionic gonadotropin (CG; #MBS263706), and prolactin (PRL; #MBS2512489) in maternal plasma were measured using rat-specific ELISA kits (MyBioSource, San Diego, CA) according to the manufacturer’s instructions. Briefly, standards and plasma samples were added to pre-coated 96-well plates in duplicates and incubated to allow antigen binding. After washing to remove unbound material, enzyme-linked detection antibodies were applied, followed by substrate solution for color development. The reaction was stopped, and absorbance was measured at 450 nm using a SpectraMax iD5 microplate reader (Molecular Devices, San Jose, CA). Concentrations were interpolated from standard curves generated using known concentrations of each analyte.

### Statistical analysis

Statistical analyses were performed in SPSS Statistics (v30, IBM, Chicago, IL, USA). Maternal traits and the outcomes from placental histology, immunohistochemical staining, and ELISA were analyzed using univariate general linear models (GLMs). For the GLM analyses, maternal exposure (saline or oxy) and maternal treatment (+/− melatonin) were included as fixed factors. Offspring traits (e.g. placental weights, fetal weights) were analyzed using generalized linear mixed models (GLMMs) to account for the nesting of the pups within their respective dams and to reduce variance within litters, with dams included as random factors and other covariates such as litter size included as random factors, where appropriate. We assessed the main effects of maternal exposure, maternal treatment, offspring sex and their respective interactions (e.g. exposure × treatment, exposure × sex). Degrees of freedom for these models were calculated using the Satterthwaite approximation. Data are presented as mean ± SEM, with dam (N) and offspring (n) sample sizes reported in each figure legend. A total of 30 dams were used across experiments, with N = 7–8 per experimental group. All figures were generated using GraphPad Prism (v10.6; GraphPad Software, San Diego, CA, USA), and statistical significance was set at p < 0.05.

## RESULTS

### Oxy exposure and melatonin treatment significantly alter maternal gestational weight gain, maternal visceral organ weights, and fetal sex distribution

Oxy exposure (p = 0.623; Fig. 1A) and melatonin treatment (p = 0.471; Fig. 1A) did not alter mean litter size at GD19.5, and neither oxy nor melatonin altered the number of resorptions at this time point (p = 1.000; p = 1.000; Fig. 1B). However, both oxy exposure (p=0.009) and melatonin treatment (p=0.008) independently altered fetal sex distribution, resulting in a significantly higher percentage of male pups compared to litters from saline-treated controls (Table 1). A significant interaction effect (p=0.011) indicated that oxy and mel in combination modestly but significantly moderated this increase. The percentage of pregestational weight gain in the period prior to mating was comparable between saline and oxy-treated dams (Fig. 1C), but at GD19.5, oxy exposure had significantly reduced maternal gestational weight gain (Fig. 1D; p<0.001). Given this reduction in maternal weight gain, we also measured individual organ weights and organ-to-body weight ratios to better understand oxy and melatonin treatment-induced changes. Oxy exposure significantly reduced maternal liver weight and liver:body weight ratio (p<0.001; Table 1), while melatonin treatment decreased small intestine weight and small intestine:body weight ratio (p=0.008; Table 1). The weights of all other maternal organs assessed were unaffected by oxy treatment or melatonin exposure (Table 1).

**Figure 1:**
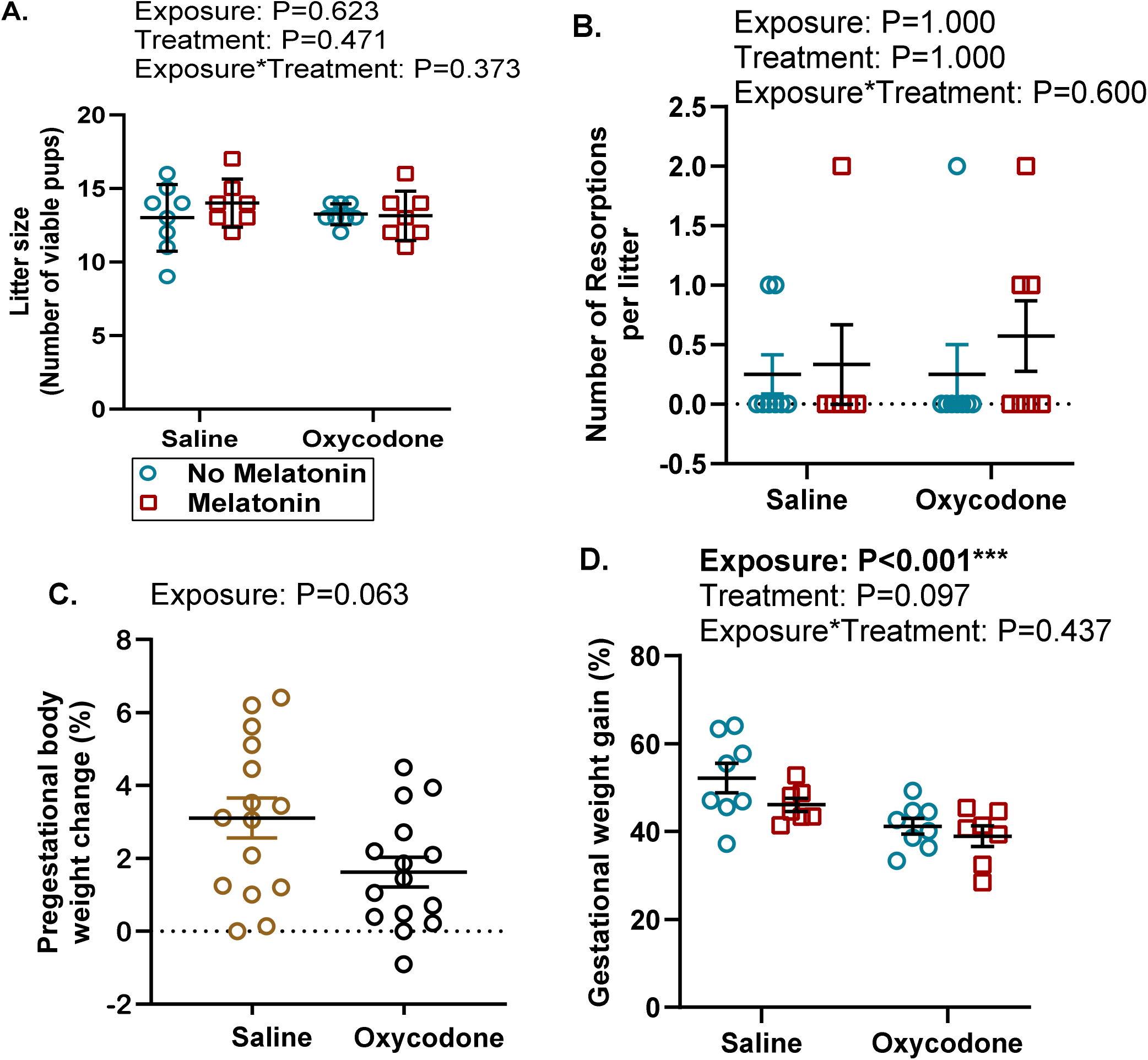
Effects of oxycodone and melatonin on litter size, resorptions, maternal anthropometry, and indices of visceral organs at GD19.5. (A) Litter size (B) Resorptions (C) Pregestational and (D) Gestational body weights. Data were analyzed by GLM and presented as mean ± SEM. N=7-8. *P*<0.05 was considered statistically significant.

**Table 1:** Effects of oxycodone and melatonin on maternal organ weights and pregnancy outcomes. Data were analyzed by GLM and presented as mean ± SEM. N=7-8. *P*<0.05 was considered statistically significant.BW denotes body weight. *P_exp_* denotes the main effect of exposure, *P_trt_* denotes the main effect of treatment, and *P_int_* denotes the main effect of interaction (exposure by treatment).

| Parameters | Saline (N=8)<br>mean±SEM | Oxy (N=8)<br>mean±SEM | Melatonin (N=7)<br>mean±SEM | OxMe (N=7)<br>mean±SEM | Statistical analysis (P-values) |
| --- | --- | --- | --- | --- | --- |
| Right Kidney weight (mg) | 1017±65.91 | 1006±26.45 | 1122±48.52 | 1007±25.23 | $P_{exp} = 0.147$<br>$P_{trt} = 0.223$<br>$P_{int} = 0.233$ |
| Right Kidney:BW ratio | 2.61±0.37 | 2.88±0.16 | 2.92±0.16 | 2.89± 0.13 | $P_{exp} = 0.321$<br>$P_{trt} = 0.212$<br>$P_{int} = 0.215$ |
| Left Kidney weight (mg) | 1083±39.54 | 1067±39.66 | 1100±44.03 | 1006±19.44 | $P_{exp} = 0.176$<br>$P_{trt} = 0.573$<br>$P_{int} = 0.332$ |
| Left Kidney:BW ratio | 2.79±0.36 | 3.05±0.34 | 2.85±0.11 | 2.88± 0.11 | $P_{exp} = 0.231$<br>$P_{trt} = 0.625$<br>$P_{int} = 0.332$ |
| Large Intestine weight (mg) | 361.13±59.21 | 346.125±64.21 | 276.7±38.22 | 287.2± 39.90 | $P_{exp} = 0.965$<br>$P_{trt} = 0.167$<br>$P_{int} = 0.802$ |
| Large Intestine: BW ratio | 0.94±0.46 | 0.99±0.47 | 0.72±0.10 | 0.82± 0.12 | $P_{exp} = 0.603$<br>$P_{trt} = 0.189$<br>$P_{int} = 0.840$ |
| Small Intestine weight (mg) | 390.2±66.4 | 400.75±56.41 | 324.29±28.87 | 231.60± 58.11 | $P_{exp} = 0.322$<br>**$P_{trt} = 0.008$**<br>$P_{int} = 0.216$ |
| Small Intestine: BW ratio | 1.00±0.16 | 1.16±0.47 | 0.84±0.34 | 0.66± 0.15 | $P_{exp} = 0.885$<br>**$P_{trt} = 0.008$**<br>$P_{int} = 0.152$ |
| Pancreas weight (mg) | 790.05±82.94 | 753.43±63.00 | 684.6±37.58 | 653.56± 39.37 | $P_{exp} = 0.634$<br>$P_{trt} = 0.156$<br>$P_{int} = 0.969$ |
| Pancreas:BW ratio | 2.03±0.61 | 2.18±0.57 | 1.78±0.28 | 1.84±0.59 | $P_{exp} = 0.600$<br>$P_{trt} = 0.140$<br>$P_{int} = 0.827$ |
| Liver weight (mg) | 9951.33±437.2 | 8395.29±403.5 | 684.6±238.9 | 653.56± 231.51 | **$P_{exp} < 0.001$**<br>$P_{trt} = 0.304$<br>$P_{int} = 0.395$ |
| Liver:BW ratio | 25.50±1.55 | 24.05±2.12 | 25.65±0.75 | 22.14±1.39 | **$P_{exp} < 0.001$**<br>$P_{trt} = 0.136$<br>$P_{int} = 0.084$ |
| Spleen weight (mg) | 649.81±41.39 | 636.01±41.92 | 682.4±24.96 | 582.6±24.34 | $P_{exp} = 0.103$<br>$P_{trt} = 0.759$<br>$P_{int} = 0.211$ |
| Spleen:BW ratio | 1.66±0.23 | 1.82±0.18 | 1.77±0.10 | 1.66±0.07 | $P_{exp} = 0.745$<br>$P_{trt} = 0.744$<br>$P_{int} = 0.115$ |
| Right Ovary weight (mg) | 123.25±8.92 | 116.31±8.70 | 119.71±10.24 | 122.0±9.80 | $P_{exp} = 0.795$<br>$P_{trt} = 0.904$<br>$P_{int} = 0.608$ |
| Right Ovary:BW ratio | 0.32±0.02 | 0.34±0.03 | 0.31±0.03 | 0.35±0.02 | $P_{exp} = 0.306$<br>$P_{trt} = 0.897$<br>$P_{int} = 0.761$ |
| Left Ovary weight (mg) | 110.86±11.53 | 106.31±5.66 | 108.33±6.25 | 114.5±3.92 | $P_{exp} = 0.909$<br>$P_{trt} = 0.689$<br>$P_{int} = 0.450$ |
| Left Ovary:BW ratio | 0.29±0.09 | 0.31±0.04 | 0.28±0.04 | 0.33±0.03 | $P_{exp} = 0.147$<br>$P_{trt} = 0.736$<br>$P_{int} = 0.505$ |
| Fetal sex distribution (% males per litter) | 40.0±5.0 | 59.1±5.1 | 59.2±5.0 | 52.2±5.2 | **$P_{exp} = 0.009$**<br>**$P_{trt} = 0.008$**<br>**$P_{int} = 0.011$** |

### Melatonin co-administration partially mitigates prenatal oxy-induced fetal growth restriction

Prenatal oxy exposure significantly reduced mean placental weight (p<0.001; Fig. 2A), fetal weight (p<0.001; Fig. 2B), and the fetal-to-placental weight (F:P) ratio, a proxy of placental transport efficiency (p=0.011; Fig. 2C), as well as crown-rump length (p<0.001; Fig. 2D), head diameter (p=0.008; Fig. 2E), abdominal diameter (p<0.001; Fig. 2F), fetal liver weight (p<0.001; Fig. 2H), and fetal heart weight (p<0.001; Fig. 2I), with males showing greater crown-rump length crown-rump length (p<0.001; Fig. 2D) and abdominal diameter (p=0.006; Fig. 2F) than females. Melatonin treatment independently reduced mean placental weight (p<0.001; Fig. 2A), fetal weight (p<0.001; Fig. 2B), crown-rump length (p<0.001; Fig. 2D), head and abdominal diameters (p<0.001 for both; Fig. 2E and 2F), and fetal liver weight (p<0.001; Fig. 2H). However, a significant positive interaction effect indicated that the combination of melatonin and oxy significantly increased placental weight (p=0.002; Fig. 2A), fetal weight (p<0.001; Fig. 2B), crown-rump length (p<0.001; Fig. 2D), head diameter (p<0.001; Fig. 2E), and fetal heart weight (p=0.003; Fig. 2I) and also increased the head-to-abdominal ratio (p=0.017; Fig. 2G).

**Figure 2:**
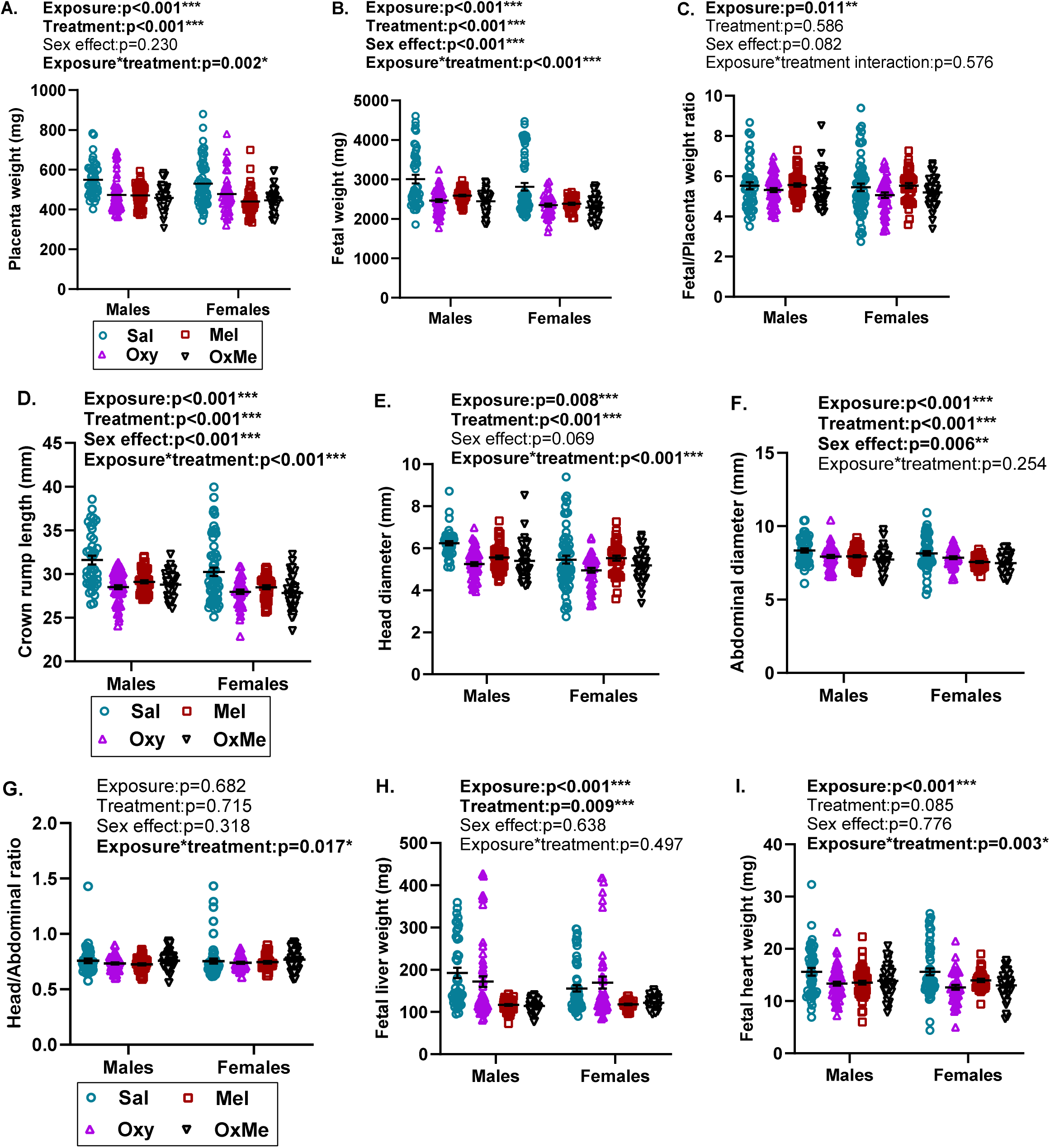
Effects of Oxy and melatonin on fetal growth and organ morphometry at GD19.5. (A) Placental weight (B) Fetal weight, (C) Fetal-to-placental weight ratio (D) Crown-rump length (E) Head diameter (F) Abdominal diameter (G) Head-Abdominal ratio (H) Fetal liver weight, and (I) Fetal heart weight. Data were analyzed by GLMM and presented as mean ± SEM. N=7-8; n=92-106/group. *P*<0.05 was considered statistically significant.

### Oxy exposure and melatonin treatment differentially alter placental structure

We assessed the impact of oxy exposure and melatonin treatment on placental zone area, as well as the cellular composition, and basal rates of proliferation and apoptosis in the labyrinth (LZ) and junctional zone (JZ). Histological analysis revealed significant expansion of the mean LZ area with prenatal oxy exposure (p= 0.025; Fig. 3A and C), which was not corrected by melatonin treatment. Neither oxy exposure nor melatonin treatment altered mean placental JZ area or LZ: JZ ratio (Fig. 3A, B, and E). Immunohistochemical analyses identified a reduction in the percentage of pan-cytokeratin-positive cells in the placental LZ with prenatal oxy exposure (p<0.001; Fig. 3F), indicating that oxy may reduce the surface area of trophoblast available for nutrient exchange and offering a potential explanation for the observed increase in LZ area. Melatonin treatment did not rescue this effect.

**Figure 3.**
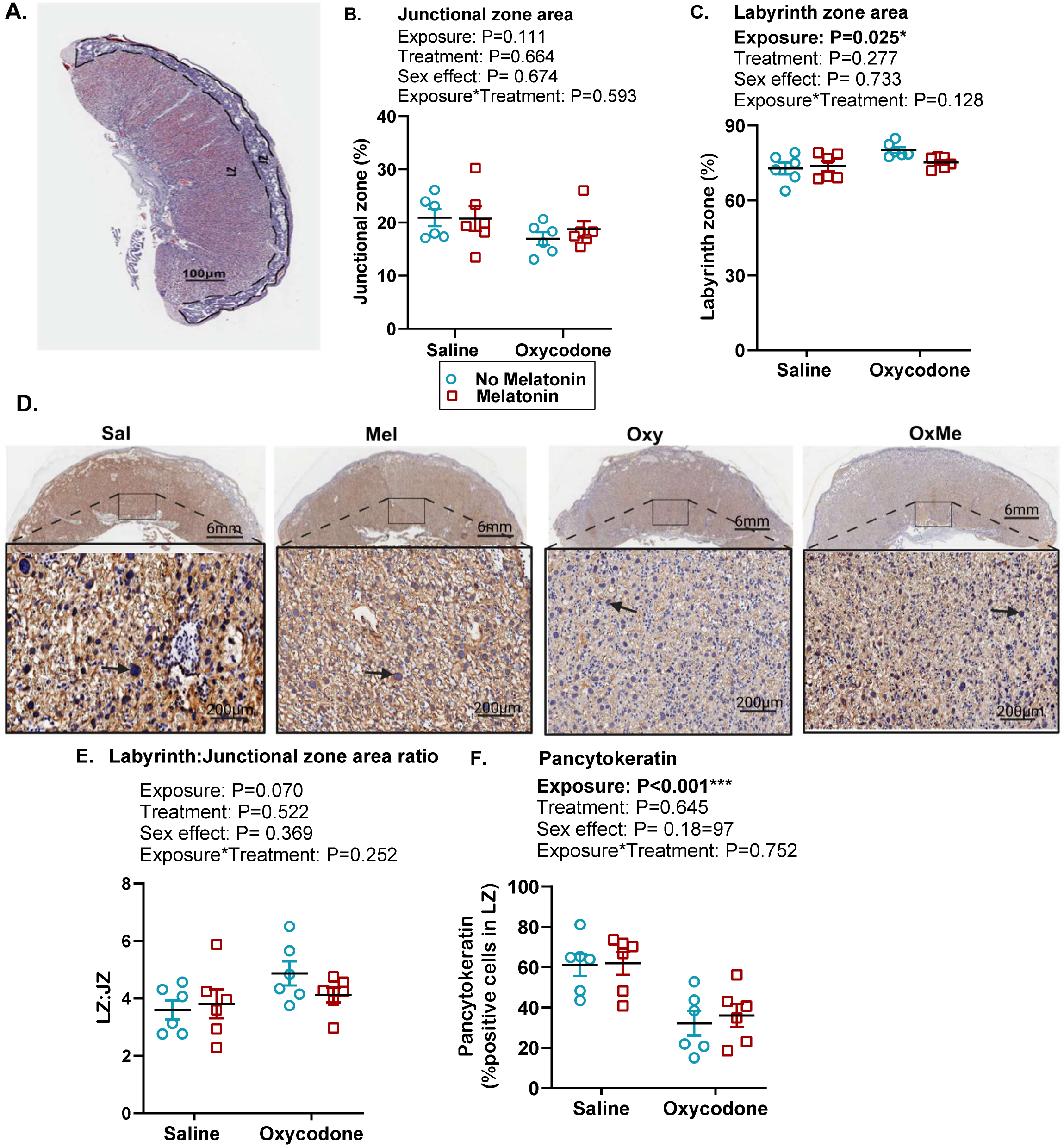
Placental zone morphometry and labyrinthine trophoblast content at GD19.5. (A) Representative H&E-stained placenta section showing delineated JZ and LZ. (B) JZ area. (C) LZ area. (D) Representative pancytokeratin immunostaining showing trophoblast cells in rat placenta. (E) LZ:JZ area ratio. (F) Quantification of pancytokeratin-positive cells in LZ. Representative images shown at 0.4X (scale bar= 6mm) and 20X (scale bar = 200 μm). Data were analyzed by GLM and presented as mean ± SEM. n=6. P<0.05 was considered statistically significant.

### Melatonin treatment significantly reduced placental proliferation and apoptosis

Melatonin treatment significantly reduced the percentage of PCNA-positive nuclei in the placental JZ (p=0.046; Fig. 4A and B), while neither oxy nor melatonin altered the percentage of PCNA-positive cells in the LZ (Fig. 4A and C). Melatonin treatment also reduced the number of TUNEL-positive cells in the JZ (Fig. 5A and B), but neither melatonin nor oxy altered the percent of TUNEL positive cells in the LZ (Fig. 5A and C). These data suggest that melatonin, but not oxy, altered basal rates of cell turnover in the placenta.

**Figure 4:**
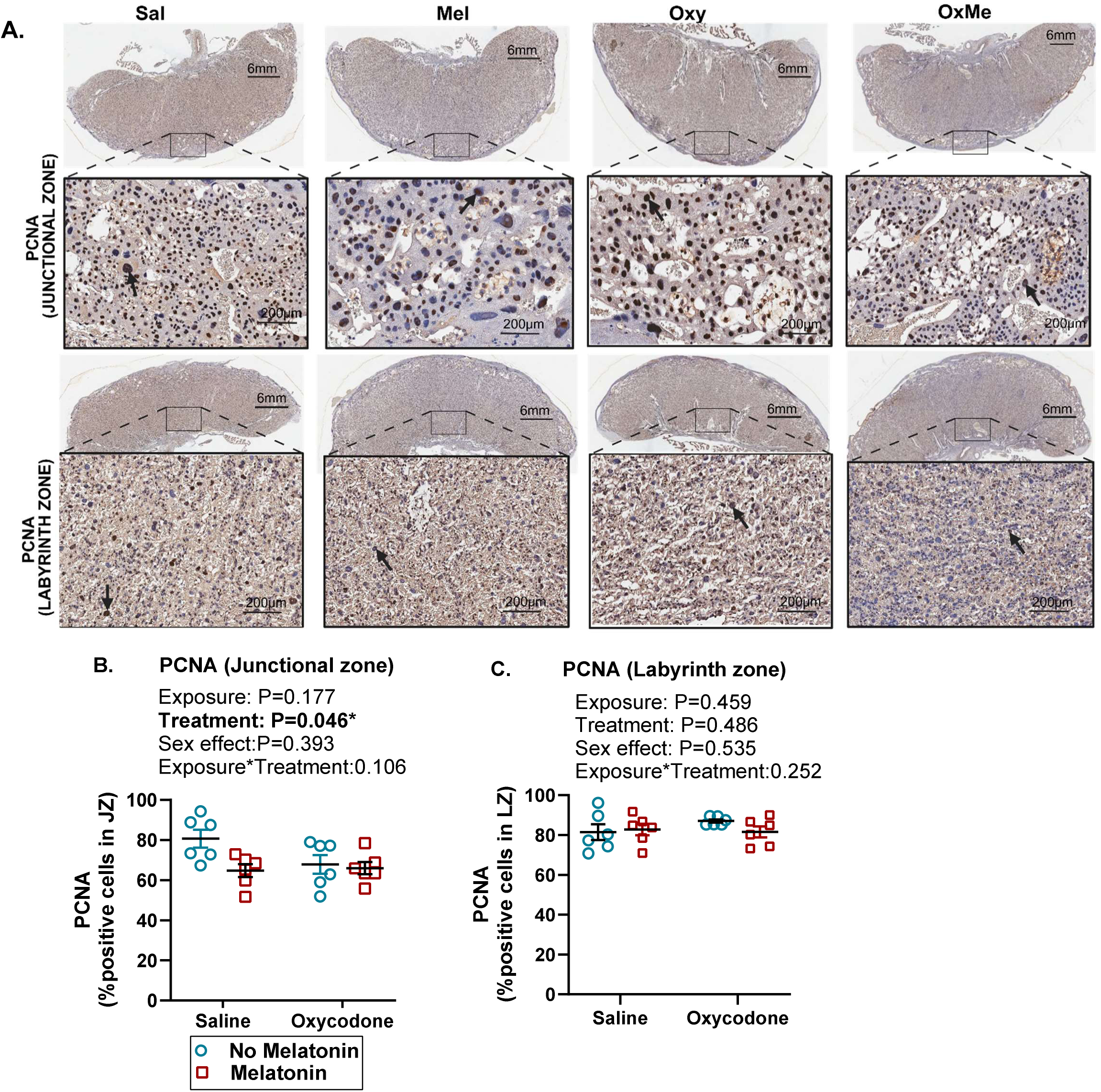
Placental cell proliferation in JZ and LZ at GD19.5. (A) PCNA immunostaining (B) Quantification of PCNA-positive cells in JZ. (C) Quantification of PCNA-positive cells in LZ. Representative images shown at 0.4X (scale bar = 6mm) and 20X (scale bar = 200 μm). Data were analyzed by GLM and presented as mean ± SEM. n=6. P<0.05 was considered statistically significant.

**Figure 5:**
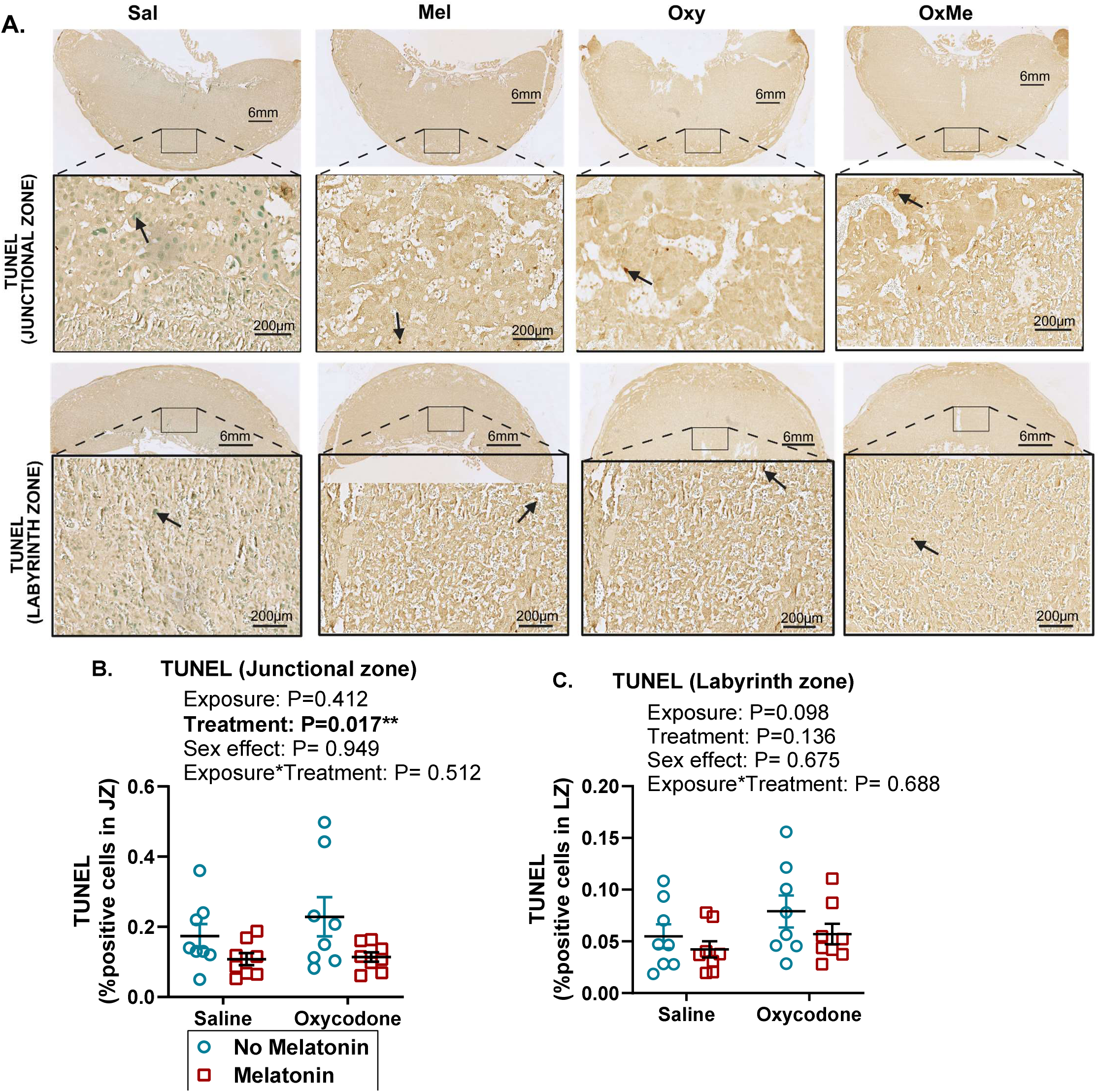
Placental cell apoptosis in JZ and LZ at GD19.5. (A) TUNEL immunostaining (B) Quantification of TUNEL-positive cells in JZ. (C) Quantification of TUNEL-positive cells in LZ. Representative images shown at 0.4X (scale bar= 6mm) and 20X (scale bar = 200 μm). Data were analyzed by GLM and presented as mean ± SEM. n=8. P<0.05 was considered statistically significant.

### Prenatal oxy exposure increased CD34 expression in the labyrinth zone

To determine whether oxy exposure or melatonin treatment affected placental vascular development and vessel maturation, we assessed the expression of alpha-SMA (α-SMA) and CD34 in the placental LZ. Neither oxy nor melatonin affected the number of α-SMA-positive cells in the LZ (Fig 6A and C). In contrast, oxy exposure significantly increased the number of CD34-positive cells in the LZ (p=0.006; Fig. 6B and D), while melatonin did not affect CD34 expression (Fig. 6B and D), suggesting an oxy-induced alteration in the composition of the placental exchange barrier.

**Figure 6:**
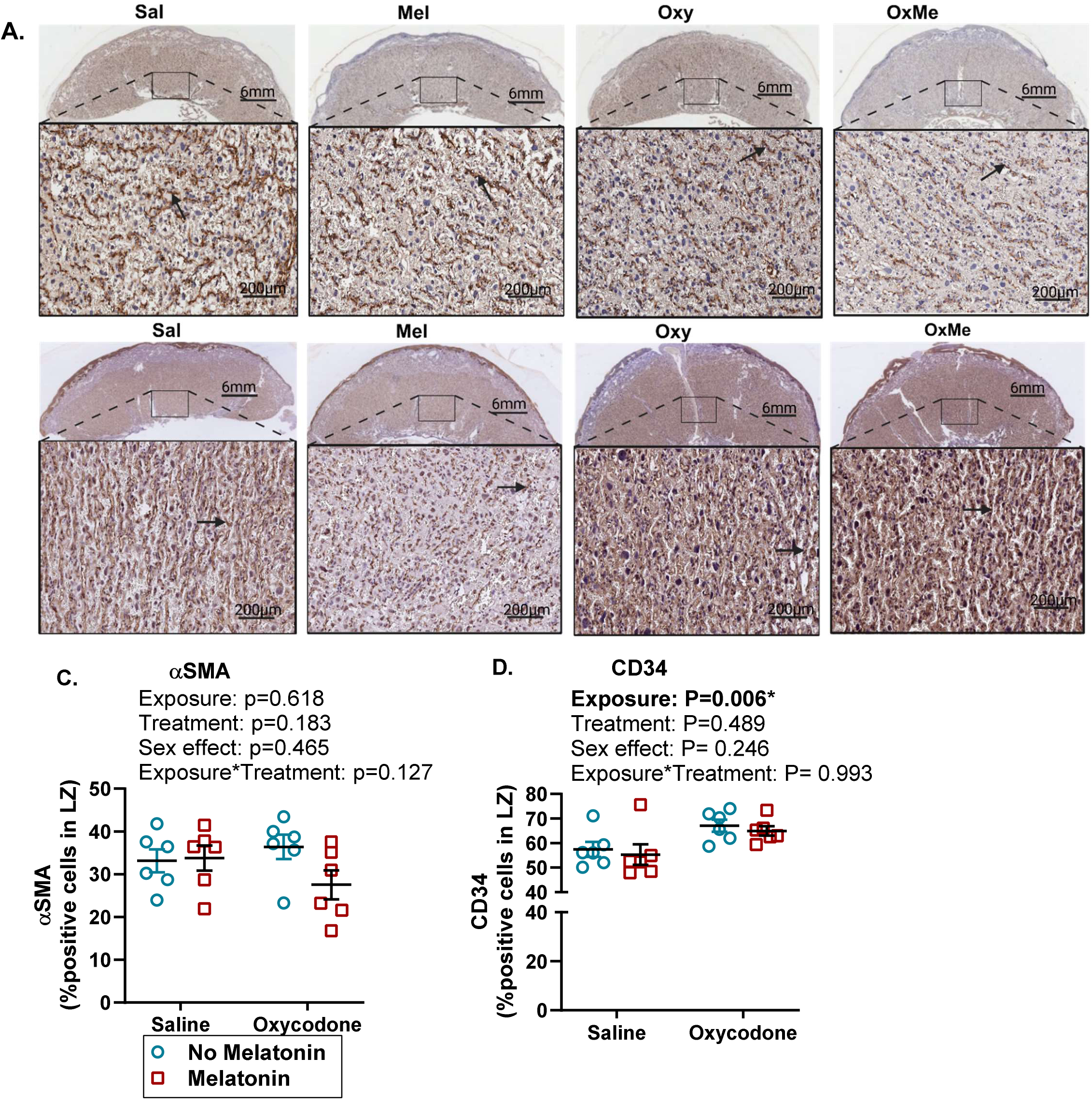
Vascular smooth muscle and endothelial cell immunostaining in the rat placental labyrinth zone. Representative images of (A) alpha-smooth muscle actin (α-SMA) and (B) CD34 immunostaining. (C) Quantification of α-SMA in LZ. (D) Quantification of CD34 in LZ. Representative images shown at 0.4X (scale bar = 6mm) and 20X (scale bar = 200 μm). Data were analyzed by GLM and presented as mean ± SEM. n=6. P<0.05 was considered statistically significant.

### Oxy exposure selectively increases maternal plasma levels of IL-1β and IL-10

To determine whether prenatal oxy exposure induced a systemic inflammatory response in the dams and their offspring, we assessed cytokine levels in the maternal and fetal plasma. Oxy exposure significantly increased the circulating concentrations of IL-1β (p<0.001; Fig. 7A) and IL-10 (p=0.003; Fig. 7F) in the maternal plasma, and melatonin treatment did not prevent these changes. Neither oxy nor melatonin affected the maternal plasma levels of any of the other cytokines (TNF-α, IL-6, CXCL1, IL-2, IL-4, and IL-5) measured in the maternal plasma and did not significantly alter cytokine levels in pooled fetal plasma (Supplementary Fig. 2).

**Figure 7:**
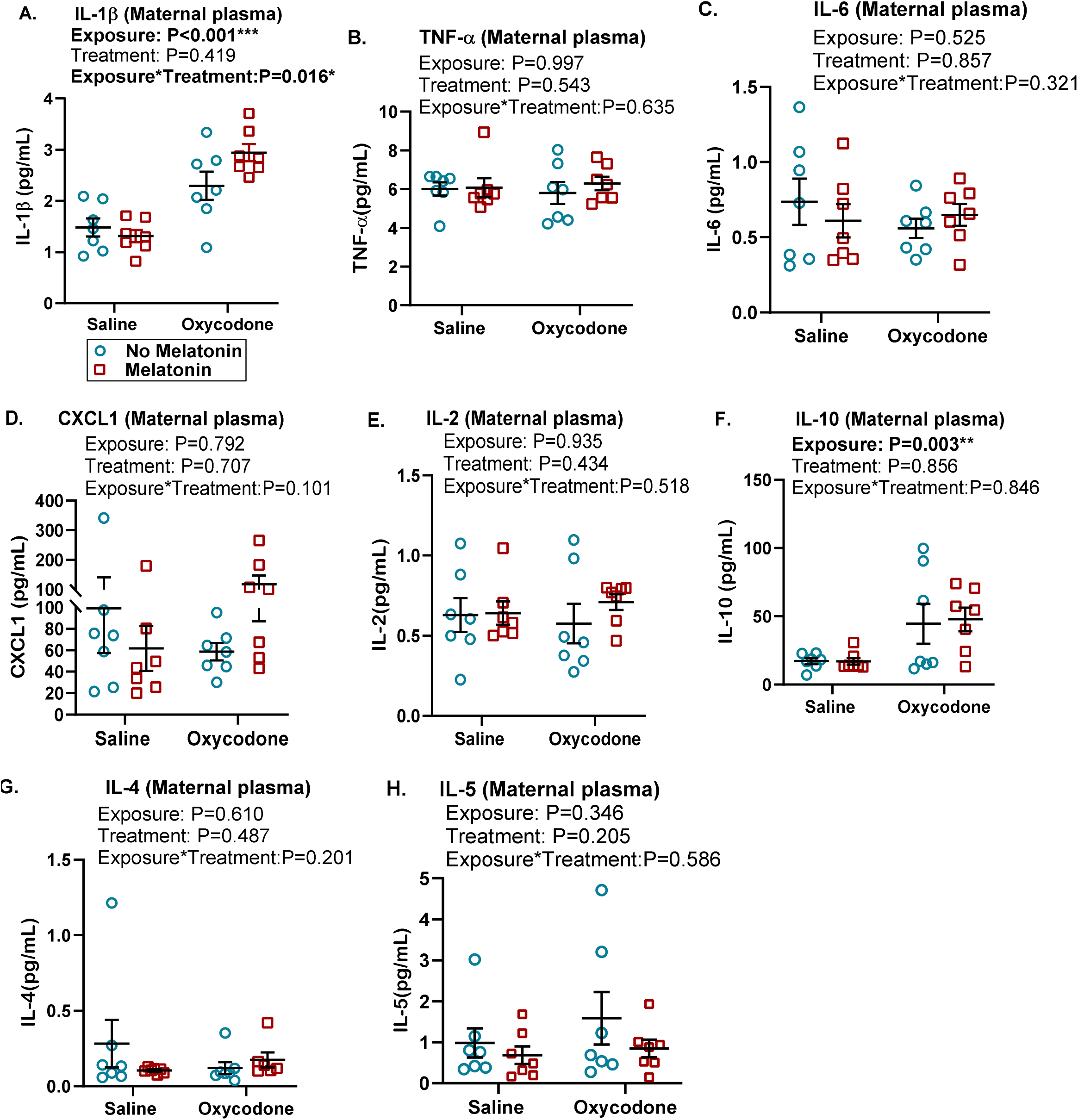
Maternal plasma cytokine profiles. **(A)** IL-1β, **(B)** TNF-α, **(C)** IL-6, **(D)** CXCL1, **(E)** IL-2, **(F)** IL-10, **(G)** IL-4, **(H)** IL-5. Data were analyzed by GLM and presented as mean ± SEM. N=7; P<0.05 was considered statistically significant.

### Oxy and melatonin treatment independently decreased maternal plasma chorionic gonadotropin levels

We assessed the levels of angiogenic factors (PlGF, VEGF, and sFlt-1) and reproductive hormones (prolactin, estriol, and CG) in the maternal plasma to determine whether oxy exposure or melatonin treatment disrupted maternal angiogenic balance or endocrine function during pregnancy. Maternal plasma levels of PlGF, VEGF, and sFlt-1 were unaffected by either oxy or melatonin treatment (Fig. 8A-C). Similarly, prolactin and estriol levels were also unaltered (Fig. 8D, E). However, CG levels were significantly reduced by both oxy exposure (p=0.013) and melatonin treatment (p=0.006) (Fig. 8F).

**Figure 8:**
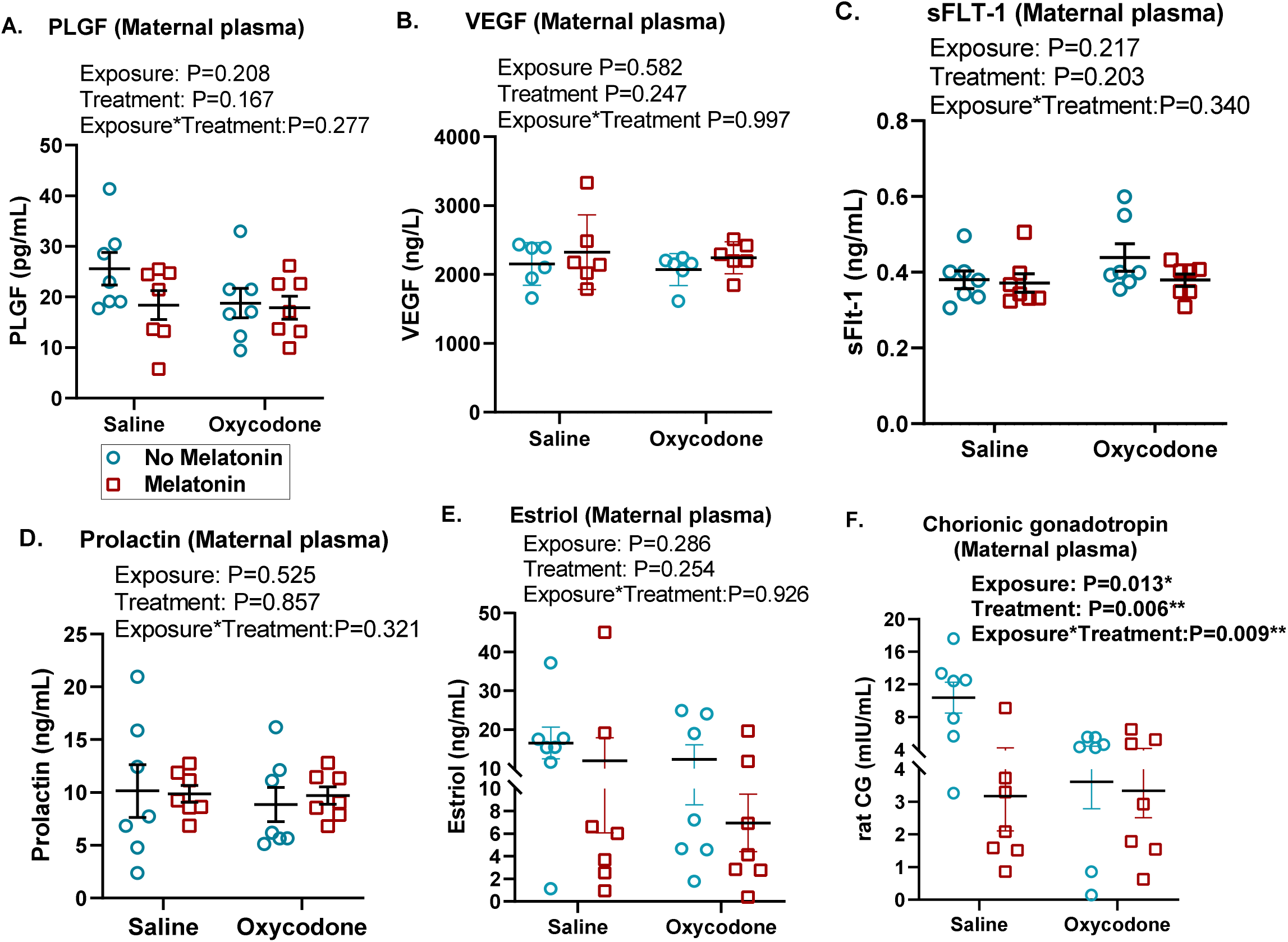
Concentrations of angiogenic factors and reproductive hormones in maternal plasma. **(A)** PLGF, **(B)** VEGF, **(C)** sFLT-1, **(D)** Prolactin, **(E)** Estriol, **(F)** rat Chorionic Gonadotropin. Data were analyzed by GLM and presented as mean ± SEM. N=7; P<0.05 was considered statistically significant.

## DISCUSSION

Previous studies investigating the effects of opioids on pregnancy outcomes have reported a range of pathologies including impaired fetal growth, placental dysfunction, altered maternal physiology and adverse pregnancy outcomes^14, 54^. It is now becoming clear that opioid exposure during pregnancy poses a significant challenge to placental function by disrupting cellular processes, vascular development, and endocrine signaling, thereby compromising overall placental integrity^26, 55, 56^. As a metabolically active organ, the placenta is particularly vulnerable to opioids, which readily cross the placental barrier and directly influence fetal development^28, 29, 57^. The extent of placental transfer varies across opioid classes, with oxy demonstrating the highest transfer rates across human trophoblast cell monolayers, raising significant concern about its impact on the placenta and fetus during gestation^28^. However, the effects of prenatal oxy exposure have not yet been fully characterized and the underlying mechanisms which lead to placental dysfunction and adverse fetal outcomes have not yet been fully explored. Therefore, the present study sought to investigate the impact of prenatal oxy exposure on pregnancy outcomes and placental biology in a rat model, while also evaluating the potential of the pleotropic hormone melatonin in attenuating oxy-induced adverse effects.

The present study showed significant reductions in gestational weight gain by GD19.5 in the oxy exposed group, with a concomitant decrease in liver weight, suggesting that oxy exposure may be causing a systemic metabolic dysregulation or altered nutritional status in the dam. The maternal effects likely compound the adverse effects of the drug on the offspring^58^, as evidenced by the reduced placental and fetal weights in the offspring of oxy exposed dams. The absence of an effect of oxy or melatonin on mean litter size and resorption rate indicates that neither intervention has a significant effect on ovulation or embryo implantation and are compatible with pregnancy success. However, the observed reduction in fetal and placental weights following oxy exposure is consistent with several studies that reported FGR in pregnancies complicated by opioid use disorder ^13, 59^. Additionally, the decrease in the F:P ratio indicates a decline in placental efficiency, meaning that for every gram of placental tissue, less fetal growth is supported^60^. This reduction in efficiency often reflects a failure of the placenta to meet the metabolic demands of the developing fetus, a hallmark of inadequate nutrient and oxygen transfer across the maternal-fetal interface ^17, 60^. In our rat model, oxy-induced FGR was accompanied by reductions in other morphometrics, including crown-rump length, head diameter, and abdominal diameter, with evident sex-specific differences as males exhibit greater crown-rump length and abdominal diameter than females despite the FGR. This is consistent with previous studies that reported that males generally display larger body dimensions and weights compared to females, even under growth-restricted conditions^61, 62^ and this sexual dimorphism suggests that male offspring prioritize structural growth more than females when subjected to intrauterine challenges^61–64^. Additionally, the significant decrease in fetal liver and heart weights in oxy-exposed fetuses provides further supportive evidence for oxy-induced FGR; for instance, fetal liver development is particularly sensitive to maternal nutrient availability and placental function^65, 66^; thus, its reduced mass may reflect diverted resources or direct effects of oxy on hepatocyte proliferation or metabolism.

While melatonin treatment is generally considered safe and beneficial, we observed that melatonin independently reduced fetal and placental weights and some growth parameters in our rat model, but, when combined with oxy, it partially improved some fetal outcomes including placental and fetal weights, crown rump length, head diameters and the head/abdominal ratio indicative of a potential brain sparing effect. This is an adaptive fetal response usually triggered by an adverse in utero environment conditions, to prioritize blood flow to protect the development of the brain and heart at the expense of abdominal viscera^67–69^. The paradoxical effects of melatonin on fetal growth in pre-clinical models are not uncommon, for instance, some studies have shown fetal weight increases, while others in hypoxic or growth-restricted models report inconsistent or even negative effects on growth metrics, and all of these have been attributed to melatonin’s actions on cellular proliferation, circadian entrainment, and metabolic regulation^70–73^. Accumulating evidence suggests that the divergent outcomes of melatonin administration are context-dependent, whereby melatonin’s actions are shaped by the underlying intrauterine environment^70, 71, 74^. In compromised pregnancies, such as FGR or hypoxia, melatonin appears to exert adaptive, cytoprotective effects that can support placental function and fetal organ development^75–77^. However, under physiological conditions, melatonin may modulate the developmental processes, leading to adverse changes in growth trajectories rather than giving beneficial effects^71, 73, 74^. Collectively, the findings of our present study show that the interaction between oxy and melatonin causes partial recovery of some growth parameters, although not to control levels, and this indicates that the detrimental effects of oxy are not fully reversible by melatonin.

Placental structural and cellular alterations provide additional insight into the mechanisms underlying impaired fetal growth in our model. Histological analysis revealed that prenatal oxy exposure caused an expansion of the LZ. In the rodents, the LZ is the primary site of nutrient and gas exchange between maternal and fetal circulations. An expansion of this zone in the context of reduced placental weight and efficiency may represent a compensatory response to intrauterine stress, nutrient deficiency or hypoxia^18, 78^. By increasing the LZ area, the placenta may be attempting to maximize exchange surface area to counter oxy-mediated disruptions. However, this adaptation was unsuccessful, given that FGR was observed in these animals. Our observation of a concomitant reduction in the percentage of pan-cytokeratin-positive cells in the LZ of oxy-exposed animals provides evidence of oxy-induced dysregulation of placental development, which would likely result in a reduction in the surface area of the trophoblast exchange barrier. This may be one reason for the observed expansion of the LZ, although further stereological, transcriptomic and functional studies will be required to make definitive conclusions on the causes and consequences of this observation. Similar dissociations between structural expansion and functional impairment have been reported in rodent models of placental dysfunction, where compensatory growth does not necessarily translate into improved transport efficiency^18, 79^.

Prenatal oxy exposure also increased the percent of CD34-positive cells in the LZ without affecting the percent of α-SMA-positive cells, a marker of more mature and stabilized vessels. CD34 is widely used as a marker of endothelial cells and its upregulation is commonly associated with angiogenesis and increased microvascular density^80, 81^. The selective increase in CD34 expression, in the absence of a corresponding change in α-SMA, suggests an expansion of immature or poorly stabilized vascular structures^82^. This pattern may reflect a compensatory angiogenic response to intrauterine stress or impaired nutrient delivery induced by oxy exposure, a mechanism that has previously been described in conditions of placental insufficiency^81, 83^. Consistent with this interpretation, our findings support a compensatory angiogenic response in which increased vessel formation is not accompanied by an increase in mature vasculature in the LZ.

While prenatal oxy exposure primarily affected the LZ, melatonin treatment primarily influenced the JZ, where it significantly reduced basal rates of proliferation and apoptosis. The JZ is responsible for endocrine production and structural support, and the effect of melatonin on this zone suggests that it may be promoting a more quiescent, stable cellular state which reflects a general suppression of cellular turnover, rather than a shift toward either expansion or degeneration.

Given that maternal opioid exposure has been reported to trigger several inflammatory signaling pathways leading to excessive release of proinflammatory cytokines^84–86^, we investigated whether prenatal oxy exposure would induce a systemic inflammatory response in the dams and their offspring, and if melatonin treatment would mitigate oxy-induced systemic inflammation. We found that oxy exposure induced a targeted inflammatory response rather than triggering systemic inflammatory cytokine storm, by selectively increasing maternal plasma concentrations of IL-1β and IL-10; melatonin treatment did not attenuate these pro-inflammatory responses. Opioids are known to act via Toll-like receptors and mu-opioid receptors on immune cells, promoting the release of IL-1β^84, 87, 88^. Opioids can also signal via Toll-like receptor-4 on human and murine trophoblast lineages, resulting in cytokine release and an inflammatory profile that can impair nutrient transport and cellular communication^56, 88–90^. The elevation of maternal IL-10 that we observed may represent a compensatory maternal response aimed at dampening this opioid-induced inflammation. Interestingly, these cytokine changes were not observed in the fetal plasma, suggesting that the maternal systemic inflammation is not transmitted to the fetus, or that the fetal immune system responds via different mechanisms not captured in the fetal circulation. Nevertheless, maternal systemic inflammation can indirectly affect the fetus by altering placental perfusion or nutrient transporter expression, thus contributing to the observed growth restriction^63, 91^. Overall, our findings are consistent with the concept that maternal inflammation can influence placental function without necessarily inducing parallel changes in the fetal compartment^92^.

The independent reduction in maternal plasma CG by both oxy and melatonin represents another crucial dimension of our findings in this present study. CG supports corpus luteum function and progesterone production during early pregnancy and has critical roles in uterine and placental vascular remodeling^93^. In rats, where placental luteotropic function is maintained in part through analogous gonadotropin signaling, the reduced circulating levels provide more evidence of JZ dysfunction and impaired endocrine output, as JZ trophoblast giant cells are the primary source of CG. Reduced gonadotropin levels may also impair the placenta’s ability to signal maternal recognition of pregnancy or modulate the maternal immune system, potentially creating a less hospitable environment for fetal development^26, 93^. Melatonin also reduced plasma CG levels in our rat model, suggesting that exogenous melatonin may influence the maternal endocrine axis regardless of its antioxidant roles. Interestingly, other endocrine markers, including prolactin and estriol, as well as angiogenic factors such as VEGF, PlGF, and sFlt-1, were not altered by any intervention. This selective effect suggests that oxy and melatonin do not broadly disrupt endocrine signaling, but rather they target specific hormonal pathways.

We acknowledge that our present study has some limitations that require further investigations. While we observed FGR and alterations in vascular markers were identified, functional assessments of blood flow or nutrient transport were not performed. Also, while we observed changes in maternal plasma cytokine levels, the lack of tissue-specific cytokine analysis in the placenta and fetal brain limits our ability to map the exact pathways of neuro-inflammatory transfer. Additionally, while the rodent model is an excellent model for studying placental structure and function, the inherent differences in CG-like hormone signaling compared to human CG, necessitates caution in direct clinical translation of our findings.

Despite these limitations, our findings show that prenatal oxy exposure disrupts placental structure, labyrinth anatomy and endocrine function, and induces maternal systemic inflammation, leading to significant FGR. The expansion of the placental LZ and increased expression of markers for immature vessels suggest a compensatory but ineffective response to oxy. Melatonin co-administration provides a partial rescue of fetal and placental growth and facilitates a brain-sparing response, highlighting its potential as a neuroprotective agent in the context of prenatal oxy exposure. Overall, our study identifies the placenta as a critical mediator of opioid-induced developmental harm and emphasizes the importance of monitoring placental health and maternal inflammatory and endocrine status in pregnancies complicated by opioid use. Future research requires a closer focus on the developmental trajectories of individual cell types in oxy-exposed placentas, as well as in depth analyses of nutrient partitioning, and long-term neurodevelopmental outcomes of exposed fetuses to determine if the brain sparing response induced by melatonin co-administration is truly adaptive and translates to functional neurological preservation.

## Author Contributions

**IOA** - Data curation, Formal analysis, Investigation, Project administration, Validation, Visualization, Writing – original draft, Writing – review & editing. **HMK** - Data curation, Formal analysis, Investigation, Project administration, Validation, Visualization, Writing – original draft, Writing – review & editing. **PNM** - Investigation, Validation, Writing – review & editing. **COA** - Investigation, Validation, Writing – review & editing. **VLS** - Investigation, Methodology, Project administration, Writing – review & editing. **JSD** - Conceptualization, Funding acquisition, Resources, Writing – review & editing. **ESP** - Conceptualization, Funding acquisition, Resources, Writing – review & editing. **GP** - Conceptualization, Funding acquisition, Methodology, Resources, Supervision, Writing – review & editing. **LKH** - Conceptualization, Funding acquisition, Investigation, Methodology, Project administration, Resources, Supervision, Writing – original draft, Writing – review & editing.

---

**Supplementary Figure 1:**
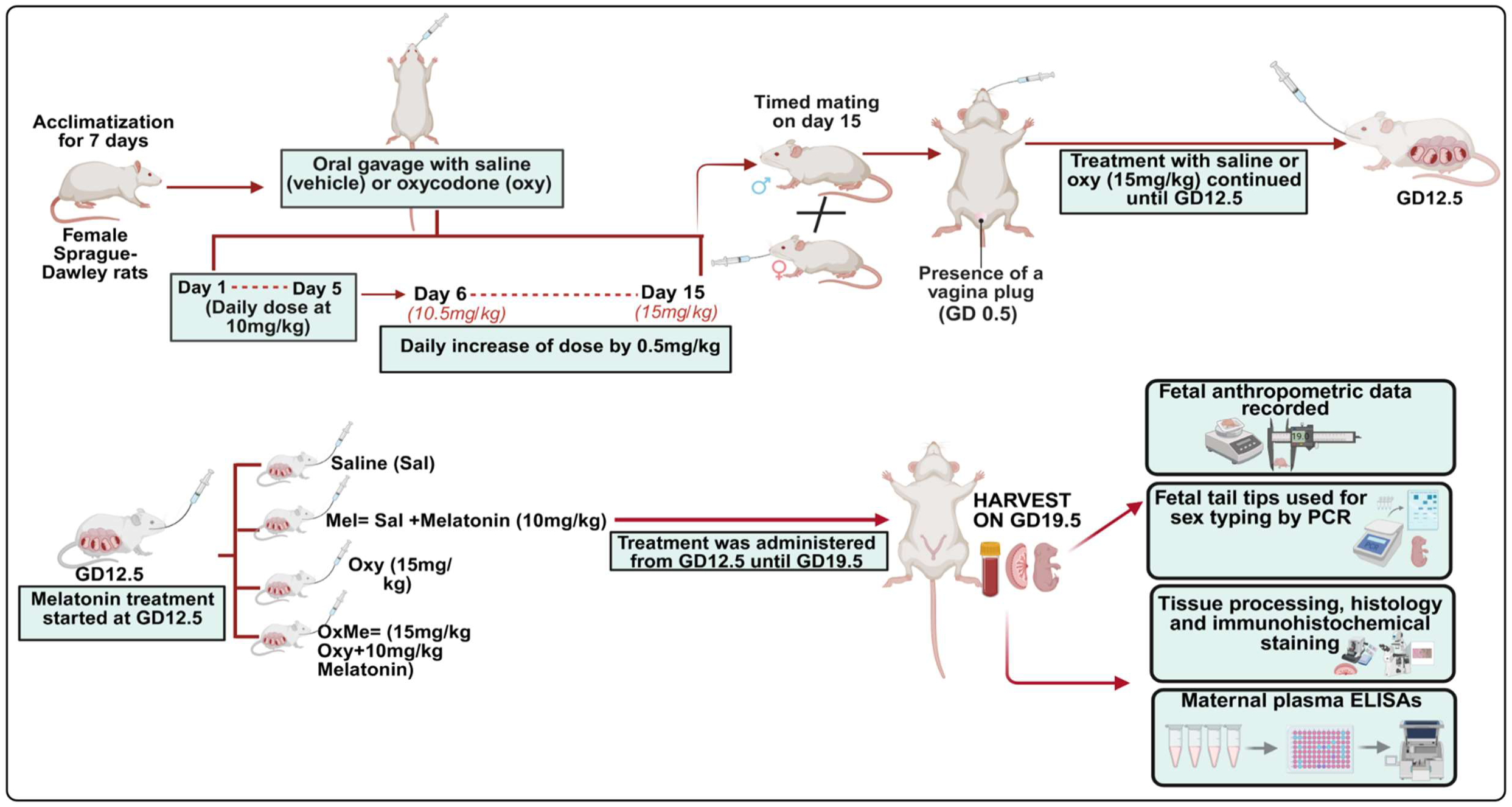
Overview of the study design.

**Supplementary Figure 2:**
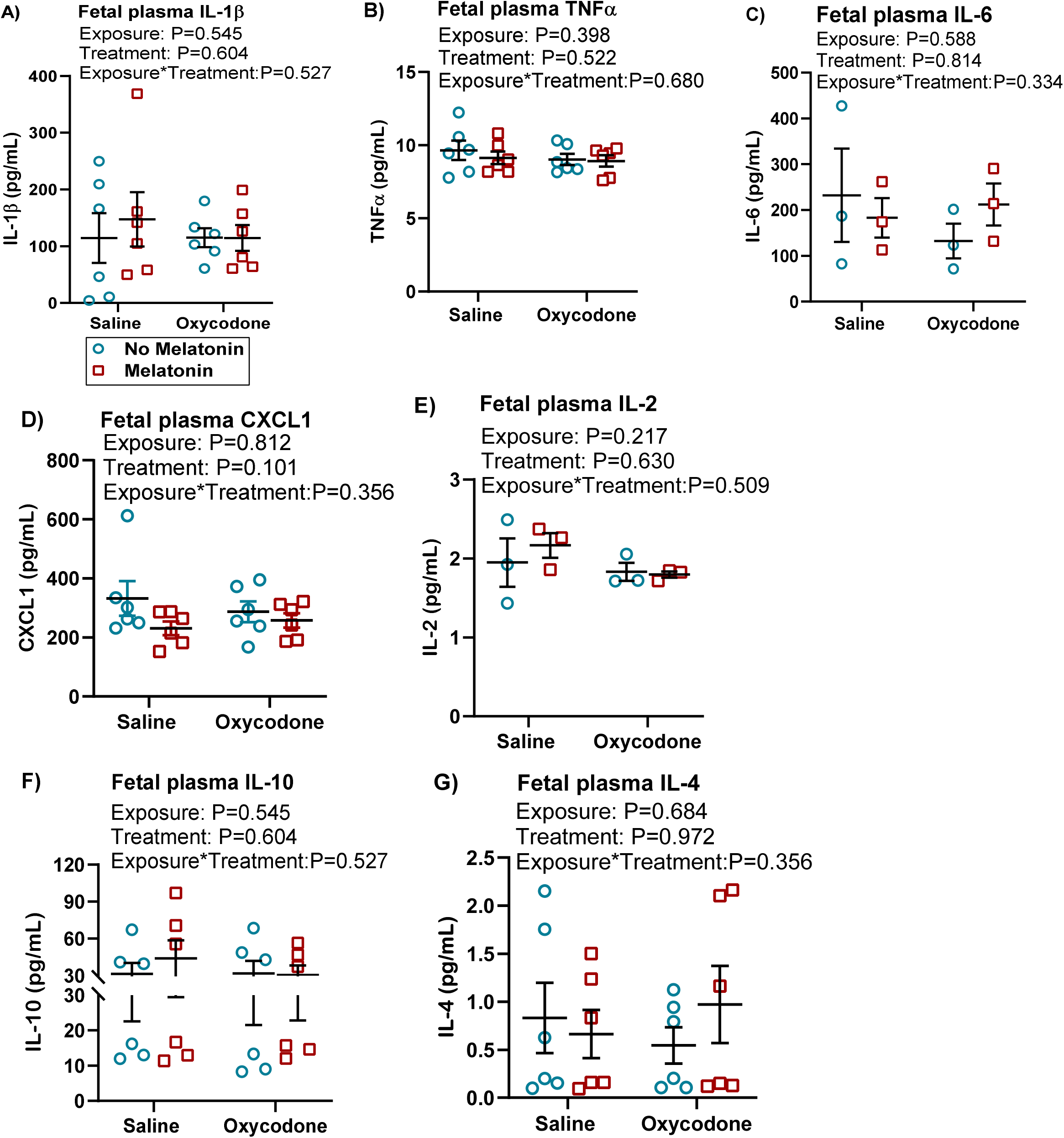
Fetal plasma cytokine concentrations. (A) IL-1β (B) TNF-α (C) IL-6 (D) CXCL1 (E) IL-2, (F) IL-10 (G) IL-4. Data were analyzed by GLM and presented as mean ± SEM. n=3-6; P<0.05 was considered statistically significant.

